# The divergence history of European blue mussel species reconstructed from Approximate Bayesian Computation: the effects of sequencing techniques and sampling strategies: Evaluating spectrum-based ABC inference in mussels

**DOI:** 10.1101/259135

**Authors:** Christelle Fraïsse, Camille Roux, Pierre-Alexandre Gagnaire, Jonathan Romiguier, Nicolas Faivre, John J. Welch, Nicolas Bierne

## Abstract

Genome-scale diversity data are increasingly available in a variety of biological systems, and can be used to reconstruct the past evolutionary history of species divergence. However, extracting the full demographic information from these data is not trivial, and requires inferential methods that account for the diversity of coalescent histories throughout the genome. Here, we evaluate the potential and limitations of one such approach. We reexamine a well-known system of mussel sister species, using the joint site frequency spectrum (jSFS) of synonymous mutations computed either from exome capture or RNA-seq, in an Approximate Bayesian Computation (ABC) framework. We first assess the best sampling strategy (number of: individuals, loci, and bins in the jSFS), and show that model selection is robust to variation in the number of individuals and loci. In contrast, different binning choices when summarizing the joint site frequency spectrum, strongly affect the results: including classes of low and high frequency shared polymorphisms can more effectively reveal recent migration events. We then take advantage of the flexibility of ABC to compare more realistic models of speciation, including variation in migration rates through time (i.e. periodic connectivity) and across genes (i.e. genome-wide heterogeneity in migration rates). We show that these models were consistently selected as the most probable, suggesting that mussels have experienced a complex history of gene flow during divergence and that the species boundary is semi-permeable. Our work provides a comprehensive evaluation of ABC demographic inference in mussels based on the coding site frequency spectrum, and supplies guidelines for employing different sequencing techniques and sampling strategies. We emphasize, perhaps surprisingly, that inferences are less limited by the volume of data, than by the way in which they are analyzed.

## Introduction

The biodiversity we inherited from the Quaternary was shaped by the process of species formation (Hewitt 2000). A long-standing question concerns the timing and rate of gene exchange that occurred while populations diverged, during the incipient stages of speciation. Model-based inferences from genetic data have been used to investigate the history of gene flow (Beaumont *et al*. 2010). Special attention has been paid to the distinction between recent divergence in a strict isolation model, and older divergence with continuous migration (Nielsen & Wakeley 2001), although of course, more complex scenarios are also possible (Marino *et al*. 2013, Sousa & Hey 2013).

With next-generation sequencing technologies, thousands of SNPs throughout the genome can be used to infer the demographic histories of non-model species pairs (Sousa & Hey 2013). One way of summarizing the information in these data is the unfolded joint site frequency spectrum (jSFS), i.e. the number of copies of derived alleles found in each of the two sampled species. A recent and fast maximum-likelihood method based on the jSFS (Gutenkunst *et al*. 2009) has proven useful for distinguishing continuous migration from strict isolation (e.g. in ragworts, Chapman *et al*. 2013, and beach mice, Domingues *et al*. 2012). The method can also evaluate more complex scenarios (e.g., in sea bass, Tine *et al*. 2014; poplars, Christe *et al*. 2017; and whitefish, Rougeux *et al*. 2017), but it struggles to explore the parameter space in these cases. In addition, the method is not well suited for transcriptome data as model comparison by log-likelihood ratio tests assumes independence of SNPs. As a consequence simulations need to be conducted to evaluate competing models, and the computational speed advantage is lost.

As an alternative, Approximate Bayesian Computation (ABC) is a method based on simulations that avoids the need to explicitly compute the likelihood (Beaumont *et al*. 2002). As such, histories of speciation characterized both by periods of strict isolation and periods of gene exchange can easily be investigated, e.g. the scenarios of ancient migration and secondary contact. These scenarios can be extended by including two cycles of “isolation / gene exchange”, following climatic changes in the Pleistocene (Figure 1). Methods have also been developed to include genome-wide heterogeneity in migration rates (Sousa *et al*. 2013; Roux *et al*. 2013). This is consistent with the “genic view” of speciation (Wu 2001), whereby barriers to gene flow are often semi-permeable, varying in strength across the genome due to linked selection and recombination (Barton & Bengtsson 1986). A major challenge in ABC, as compared to explicit likelihood methods, is the selection of summary statistics, which involves a trade-off between loss of information and reduction of dimensionality. Several methods have been suggested to select the most appropriate statistics for a given dataset and a set of models (e.g., Wegmann *et al*. 2009; Nunes & Balding 2010; Aeschbacher *et al*. 2012); but most of these summarize the site frequency spectrum (but see, e.g. Boitard *et al*. 2016) leading to a loss of information. Recently, several ABC studies have used the site frequency spectrum directly to reconstruct the history of single populations (Boitard *et al*. 2016), or multiple populations (Xue & Hickerson 2015; Smith *et al*. 2017). However the number of statistics, i.e. the number of classes in the spectrum, increases quadratically with the number of haploid genomes sampled in the case of a two-dimensional spectrum, and even faster if more than two populations are considered. For this reason, recent studies have tested different ways of binning the site frequency spectrum to circumvent the curse of dimensionality (e.g., Smith *et al*. 2017). While accumulating new data and developing new methods, studies that evaluate the impact of the sampling strategy and the inferential method (e.g. Li & Jakobsson 2012; Robinson *et al*. 2014; Shafer *et al*. 2015; Cabrera & Palsbøll 2017; Smith *et al*. 2017) to reconstruct species divergence history will be increasingly valuable.

**Figure 1.**
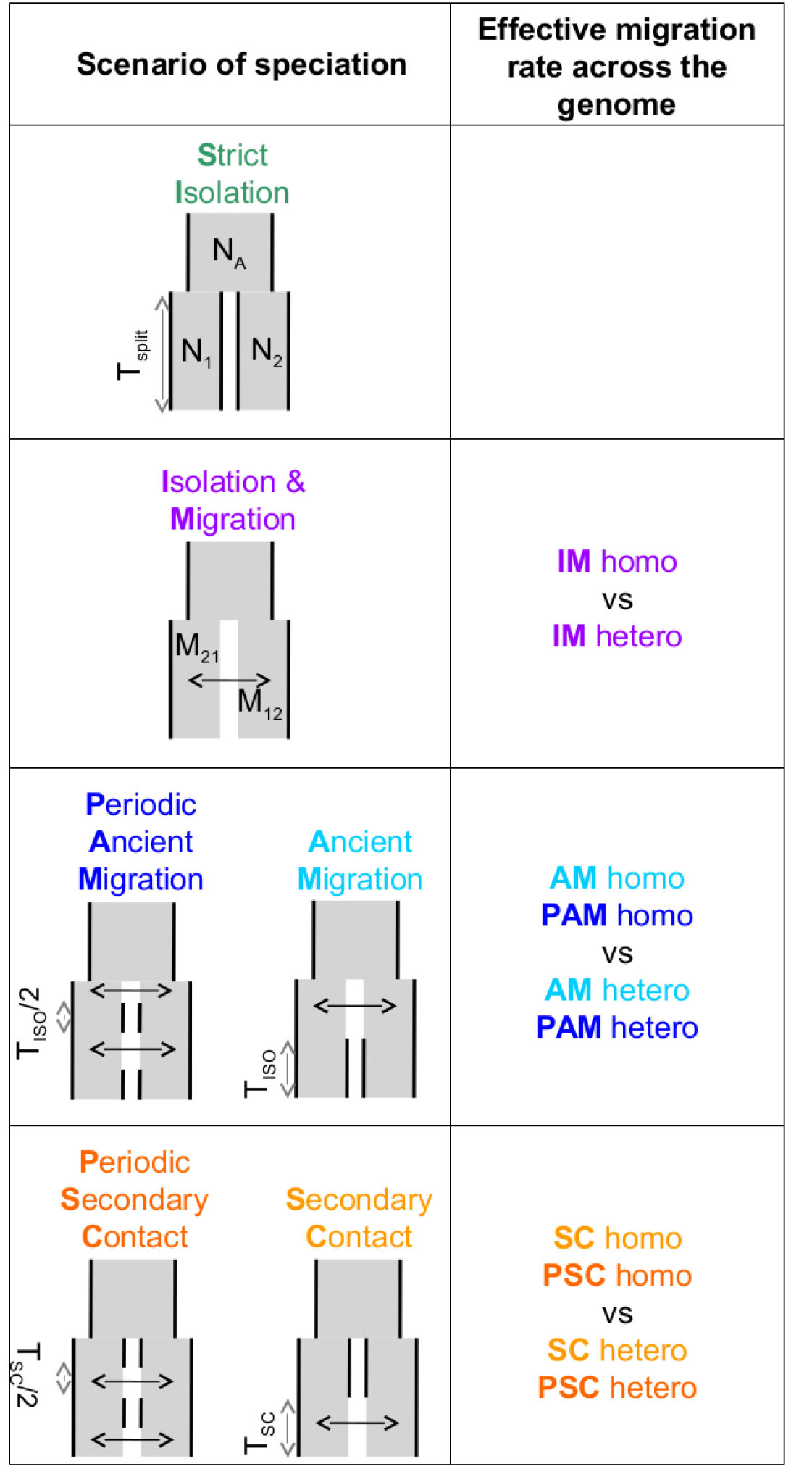
Models of speciation. Six classes of scenarios with different temporal patterns of migration are compared (left column); and for those including migration, two versions are depicted assuming either homogeneity (“homo”) or heterogeneity (“hetero”) of effective migration rate across the genome (right column). All scenarios assume that an ancestral population of effective size *N_A_* split *T_split_* generations ago into two populations of constant sizes *N_1_*, and *N_2_*. At the two extremes, divergence occurs in allopatry (SI, strict isolation) or under continuous migration (IM, isolation with migration). Through time, migration occurs at a constant rate *M_12_* from population 1 to population 2 and *M_21_* in the opposite direction. Ancient migration (AM) and periodic ancient migration (PAM) scenarios both assume that populations started diverging in the presence of gene flow. Then they experienced a single period of isolation, *T_iso_*, in the AM model while intermittent gene flow occurred in the PAM model. In the secondary contact (SC) and periodic secondary contact (PSC) scenarios, populations diverged in the absence of gene flow followed by a single period of secondary contact, *T_sc_*, in the SC model while intermittent gene flow occurred in the PSC model.

*Mytilus edulis* (Linnaeus, 1758) and *Mytilus galloprovincialis* (Lamarck, 1819) are two closely-related species that currently hybridize where their ranges overlap along the Atlantic French coasts (Bierne *et al*. 2003a) and the British Isles (Skibinski *et al*. 1983). Their interspecific barrier to gene flow is semi-permeable, and it has been shown to involve multiple isolating mechanisms, both pre-zygotic (e.g. assortative fertilization and habitat choice, Bierne *et al*. 2002, 2003b), and post-zygotic (hybrid fitness depression, Simon *et al*. 2017). Other evidence of ongoing gene flow between *M. edulis* and *M. galloprovincialis* comes from footprints of local introgression of *edulis*-derived alleles into a population of *M. galloprovincialis* enclosed within the Atlantic hybrid zone (Fraïsse *et al*. 2014). Another study (Fraïsse *et al*. 2016) revealed that the Atlantic population of *M. galloprovincialis* was more introgressed than the Mediterranean population on average. At some specific loci, however, the Mediterranean population was found to be fixed for *edulis* alleles while the Atlantic population was not introgressed at all, suggesting that an ancient contact between *M. edulis* and Mediterranean *M. galloprovincialis* occurred during glacial periods. Finally, direct model comparisons have been conducted with the IMa method of Hey & Nielsen (2007), and with an ABC framework, and shown that *M. edulis* and *M. galloprovincialis* have experienced a complex history of divergence punctuated by periods of gene flow in Europe (IMa: Boon *et al*. 2009, ABC: Roux *et al*. 2014, 2016).

Here, we analysed coding sequence datasets from this well-known pair of sister species in Europe, and systematically reconstructed its speciation history by ABC, using different sampling strategies. By different sampling strategies, we mean that we varied (i) the number of individuals sampled (2, 4, or 8), (ii) the number of SNPs (which were obtained by different sequencing techniques, “exome capture” *vs*. “rna-seq”) and (iii) the number of bins in the jSFS (binning the spectrum into 4, 7 or 23 classes). We then evaluated the influence of these choices on model selection, using eleven distinct scenarios of speciation. Our results show that the influence on inferences of the number of individuals and loci sampled is surprisingly limited, while the different ways of binning the jSFS strongly affect the results. This suggests that demographic reconstructions are nowadays more limited by the way data are analyzed than by their volume. Moreover, we find that an history of periodic connectivity, with both ancient and contemporary introgression, and a semi-permeable barrier to gene flow, best fit these data, arguing for the development of flexible inference methods to better describe complex divergence histories (Simon & Duranton 2018).

## Materials and Methods

### Sampling, sequencing, mapping and calling

Two datasets were analysed for demographic inferences. They both comprise measures of molecular polymorphism and divergence in coding sequences, obtained for the pair *M. edulis* and *M. galloprovincialis*. Although the adopted sequencing techniques were different among datasets (“exome capture” vs. “rna-seq”), the surveyed populations were similar. The usage of coding sequences to infer divergence histories of closely-related species (e.g., see Roux *et al*. 2013, 2014, 2016; McCoy *et al*. 2014; Li *et al*. 2017; Qi *et al*. 2017) is justified for several reasons: (i) synonymous mutations are less affected by direct selection than other categories of mutation, and selection affects chromosomal regions larger than genes themselves, including non-coding regions (e.g., regulatory elements, Andolfatto 2005); (ii) we implicitly modelled the effects of selection against migrant genes by including heterogeneous effective migration rates across the genome.

#### Data set 1: “exome capture”

We used the dataset already published in Fraïsse *et al*. (2016) and available on *http://www.scbi.uma.es/mytilus/index.php*. Briefly, a set of 890 EST contigs was used as a reference for a pre-capture multiplex DNA enrichment in samples of eight individuals from two geographical populations in each species (*M. edulis*: North Sea and Bay of Biscay; *M. galloprovincialis*: Brittany and Mediterranean Sea, Table 1). In addition, we used a sample of four individuals of *M. trossulus* to serve as an outgroup (Table 1). Each DNA library was sequenced twice to increase the per-base coverage (Miseq or GA2X followed by HiSeq2000). After trimming and quality-filtering, reads of each individual were aligned against the same EST reference sequences using the BWA program (bwa-mem, Li & Durbin 2009). Because of the relative divergence between the two species (~2%, Table 2), we adjusted the default parameter of BWA to allow less stringent mapping (minimum seed length *k*=10 [default: 19], clipping penalty *L*=3 [5], mismatch penalty *B*=2 [4], and gap open penalty *O*=3 [6]). Full methods are described in Fraïsse *et al*. (2016).

**Table 1.**
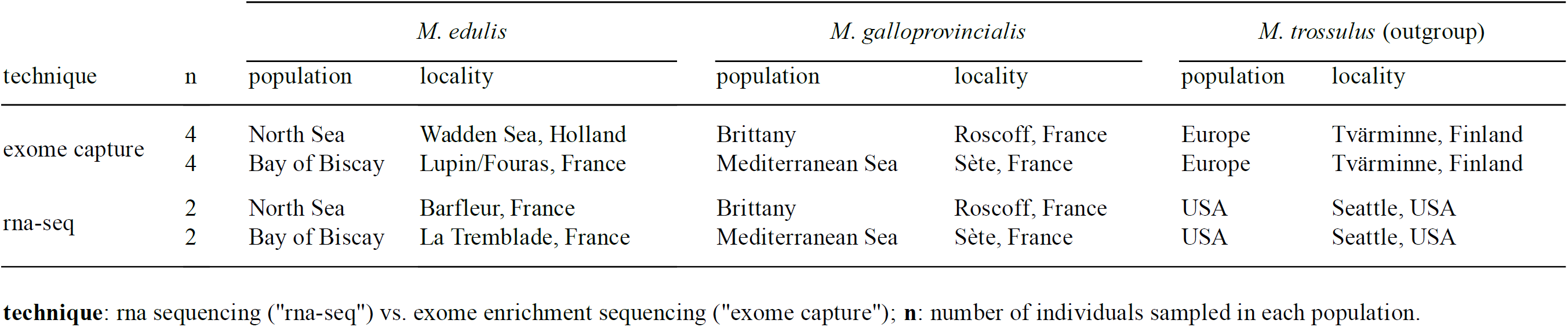
Sampling design

| technique | n | *M. edulis* |  | *M. galloprovincialis* |  | *M. trossulus* (outgroup) |  |
| --- | --- | --- | --- | --- | --- | --- | --- |
|  |  | population | locality | population | locality | population | locality |
| exome capture | 4 | North Sea | Wadden Sea, Holland | Brittany | Roscoff, France | Europe | Tvärminne, Finland |
|  | 4 | Bay of Biscay | Lupin/Fouras, France | Mediterranean Sea | Sète, France | Europe | Tvärminne, Finland |
| rna-seq | 2 | North Sea | Barfleur, France | Brittany | Roscoff, France | USA | Seattle, USA |
|  | 2 | Bay of Biscay | La Tremblade, France | Mediterranean Sea | Sète, France | USA | Seattle, USA |

**technique:** rna sequencing ("rna-seq") vs. exome enrichment sequencing ("exome capture"); **n:** number of individuals sampled in each population.

**Table 2.** Summary statistics (mscalc)

| technique | n1 | n2 | n_locus | n_SNP | S | S_sd | Sf | Sf_sd | Sx1 | Sx1_sd | Sx2 | Sx2_sd | Ss | Ss_sd |  |  |  |  |
| --- | --- | --- | --- | --- | --- | --- | --- | --- | --- | --- | --- | --- | --- | --- | --- | --- | --- | --- |
| exome capture | 2 | 2 | 516 | 3993 | 7.738 | 6.732 | 0.322 | 1.285 | 3.124 | 3.077 | 3.3 | 3.509 | 0.992 | 1.855 |  |  |  |  |
|  | 4 | 4 | 557 | 5092 | 9.142 | 8.076 | 0.097 | 0.583 | 3.555 | 3.434 | 3.896 | 4.006 | 1.594 | 2.482 |  |  |  |  |
|  | 8 | 8 | 512 | 5000 | 9.766 | 8.828 | 0.025 | 0.296 | 3.504 | 3.502 | 4.258 | 4.363 | 1.979 | 2.761 |  |  |  |  |
| rna-seq | 2 | 2 | 2147 | 17275 | 8.046 | 6.554 | 0.81 | 3.057 | 2.966 | 3.14 | 2.809 | 2.851 | 1.462 | 2.216 |  |  |  |  |
|  | 4 | 4 | 1842 | 17902 | 9.719 | 7.953 | 0.507 | 2.608 | 3.344 | 3.328 | 3.368 | 3.318 | 2.501 | 3.318 |  |  |  |  |
| | $\pi_1$ | $\pi_1\_sd$ | $\pi_2$ | $\pi_2\_sd$ | $\theta w_1$ | $\theta w_1\_sd$ | $\theta w_2$ | $\theta w_2\_sd$ | D1 | D1_sd | D2 | D2_sd | FST | FST_sd | div | div_sd | netdiv | netdiv_sd |
|  | 0.016 | 0.016 | 0.016 | 0.015 | 0.017 | 0.017 | 0.016 | 0.016 | -0.27 | 0.674 | -0.391 | 0.513 | 0.114 | 0.189 | 0.02 | 0.017 | 0.004 | 0.009 |
|  | 0.014 | 0.014 | 0.013 | 0.013 | 0.016 | 0.015 | 0.016 | 0.015 | -0.668 | 0.78 | -0.786 | 0.702 | 0.101 | 0.158 | 0.017 | 0.016 | 0.004 | 0.008 |
|  | 0.012 | 0.012 | 0.012 | 0.012 | 0.017 | 0.015 | 0.019 | 0.016 | -0.942 | 0.78 | -1.143 | 0.67 | 0.088 | 0.141 | 0.015 | 0.015 | 0.003 | 0.008 |
|  | 0.038 | 0.031 | 0.036 | 0.029 | 0.038 | 0.03 | 0.037 | 0.029 | -0.068 | 0.805 | -0.179 | 0.699 | 0.181 | 0.256 | 0.057 | 0.053 | 0.02 | 0.049 |
|  | 0.034 | 0.027 | 0.032 | 0.025 | 0.036 | 0.026 | 0.036 | 0.026 | -0.27 | 0.848 | -0.493 | 0.751 | 0.171 | 0.229 | 0.051 | 0.046 | 0.018 | 0.042 |

**technique:** rna sequencing ("rna-seq") vs. exome enrichment sequencing ("exome capture"); **n:** number of individuals analyzed in each species;
**n\_locus:** total number of locus **n\_SNP:** total number of polymorphic sites.
The following statistics were calculated for each locus. Their average (in black) and standard deviation (in grey) across all loci are given.
**S:** number of polymorphic sites; **Sf:** number of fixed differences; **Sx:** number of exclusive polymorphic sites; **Ss:** number of shared polymorphic sites;
**$\pi$ :** number of pairwise differences (Tajima, 1983); **$\theta w$ :** Watterson's $\theta$ (Watterson, 1975); **D:** Tajima's D (Tajima, 1989a, 1989b);
**FST = $1 - \pi_S / \pi_T$ :** level of species differentiation, where **$\pi_S$** is the average pairwise nucleotide diversity within species and **$\pi_T$** is the total pairwise nucleotide diversity of the pooled sample across species; **div:** total interspecific divergence; **netdiv:** net molecular divergence measured at synonymous positions.

We used a maximum-likelihood method, implemented in the program read2snps (Tsagkogeorga *et al*. 2012; Gayral *et al*. 2013), to call genotypes directly from read numbers at each position. The method computes the probability of each possible genotype after estimating the sequencing error rate. To limit bias in the site frequency estimation (Han *et al*. 2013), a minimum of 10X coverage, i.e. ten reads per position and per individual, was required to call a genotype. Only genotypes supported at 95% were retained; otherwise missing data was applied. Moreover, paralogous positions were filtered-out using a likelihood ratio test based on explicit modelling of paralogy.

#### Data set 2: “rna-seq”

The dataset 2 is made of sequenced transcriptomes (RNA-seq) previously generated in a wide meta-analysis study comparing levels of polymorphism across 76 animal species (Romiguier *et al*. 2014). It includes the transcriptomes of four individuals sampled in the same populations as described above and one individual of *M. trossulus* (Table 1). Briefly, for each individual, cDNA libraries were prepared with total RNA extracted from whole body and sequenced on HiSeq2000. Illumina reads (100 bp, paired-end) were mapped with the BWA program on *de novo* transcriptomes, independently assembled for each species with a combination of the programs Abyss and Cap3, following the strategy B and D in Cahais *et al*. (2012). Contigs with a per-individual average coverage below ×2.5 were discarded. Genotype calling was performed as described in the first data set using identical filters. Open reading frames were predicted with the Trinity package and sequences carrying no ORF longer than 200 bp were discarded. Full methods are described in Romiguier *et al*. (2014).

### Data analysis

We restricted our analysis to loci assembled in all individuals, and longer than 300 bp after filtering positions containing missing data, or more than two segregating alleles when the three species were aligned. Only synonymous positions were used. The total number of loci and SNPs retained in each dataset is given in Table 2.

#### Site Frequency Spectrum

We first computed the jSFS for each dataset (Figure 2 and Figure S1). Each derived allele, oriented by treating the outgroup sequence as a fixed ancestral allele, was assigned to one cell of the jSFS, depending on its frequency in each of the two populations. From the full spectrum, different classes of polymorphism were extracted and used as summary statistics (Tellier *et al*. 2011). Specifically, we used the four Wakeley-Hey classes (*jsfs*=4 in Figure 2): fixed differences, Sf; private polymorphisms for each species Sx_1_ and Sx_2_; and shared polymorphisms, Ss (Wakeley & Hey 1997). We also considered a summary in which the Sf and Sx classes were split depending on whether the derived allele was fixed or absent in the other species (*jsfs*=7 in Figure 2; Ramos-Onsins *et al*. 2004). The third decomposition of the jSFS contains twenty-three classes of polymorphisms because singletons and doubletons in each population were included as new classes (*jsfs*=23 in Figure 2). This corresponds to the full spectrum with *n*=2 diploid individuals.

**Figure 2.**
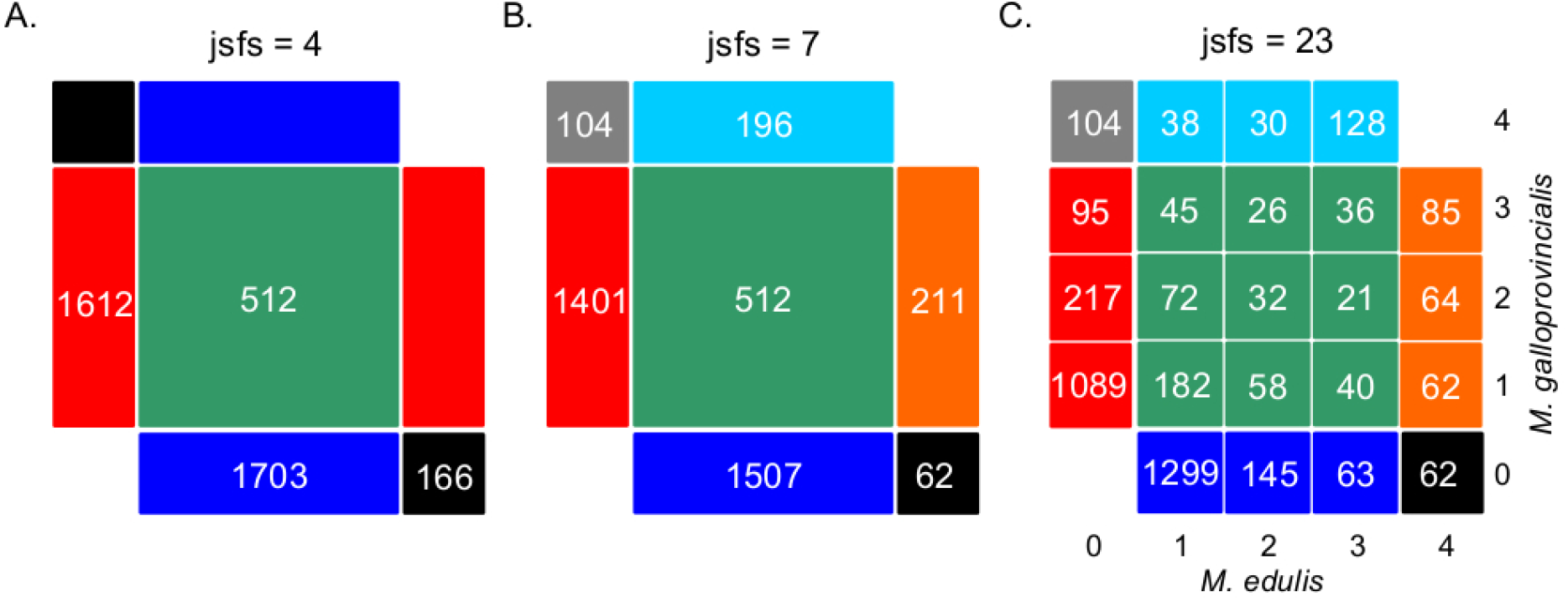
Decomposition of the unfolded joint site frequency spectrum for *n*=2 individuals (i.e. 4 alleles) in each species. The density of derived alleles in species 1 (M *edulis*, x axis) and species 2 *(M. galloprovincialis*, y axis) is indicated by a number within each cell. Only sites showing two distinct alleles in the inter-specific alignment were considered, hence the cells {0;0} and {4;4} have been masked. The total number of polymorphic sites is 3993 SNPs (“exome capture” data). (A) Decomposition of the jSFS into four classes of polymorphism without an outgroup sequence (i.e., the Wakeley-Hey classes): fixed differences (black), private polymorphisms in species 1 (blue) or species 2 (red) and shared polymorphisms (green). (B) Decomposition of the jSFS into seven classes of polymorphism by using the sequenced outgroup. Two alleles are differentially fixed between the two species: the derived allele can be fixed in species 1 (black) or in species 2 (grey). Exclusive polymorphism can be the result of a recent mutation specific to species 1 (blue) or species 2 (red); but it can also be the result of an ancestral mutation only fixed in species 2 (cyan) or in species 1 (orange). Shared poly morph isms are shown in green. (C) Decomposition of jSFS into twenty-three classes of polymorphism. Singletons and doubletons in each species were included as new classes. Note that in the case of *n*=2, this is the full spectrum.

#### Estimators of polymorphism and divergence

We then computed a set of genetic statistics across loci to make use of the coalescent information contained within each sequence, instead of considering each SNP separately. Following previous studies (e.g., Fagundes *et al*. 2007; Ross-Ibarra *et al*. 2008; Roux *et al*. 2011), we used the following statistics: (1) nucleotide diversity, π_1_ and π_2_ (Tajima 1983); (2) Watterson’s θ_W1_ and θ_W2_ (Watterson 1975); (3) total and net interspecific divergence, div and netdiv; (4) between-species differentiation, FST, computed as 1-π_S_/π_T_, where π_S_ is the average pairwise nucleotide diversity within species and π_T_ is the total pairwise nucleotide diversity of the pooled sample across species. We also included the four Wakeley-Hey’s classes as explained above (Sf, Sx_1_, Sx_2_ and Ss). Finally, we assessed departure from mutation/drift equilibrium using Tajima’s D_1_ and D_2_ (Tajima 1989a,b). The average and standard deviation across the loci of these statistics were calculated with the program MScalc (available from *http://www.abcgwh.sitew.ch/*, see Roux *et al*. 2011), and their values are given in Table 2 (“mscalc”).

### Inferences by Approximate Bayesian Computation (ABC)

#### Scenarios of speciation

Six distinct scenarios of speciation were considered (Figure 1). Each scenario modeled an instantaneous division (occurring T_split_ generations ago) of the ancestral population of effective size N_A_ into two populations of constant sizes N_1_ and N_2_. The Strict Isolation scenario (SI) assumed that divergence occurred without gene exchange between the two populations. The other models differed by their temporal pattern of migration, which occurred at a rate M_12_ from population 1 to population 2, and a rate M_21_ in the opposite direction. Ancient Migration (AM) and Periodic Ancient Migration (PAM) scenarios both assumed that migration was restricted to the early period of divergence. In the AM scenario, the two populations experienced a single period of strict isolation (T_iso_) while in the PAM scenario, migration was stopped twice with an intermediate period of isolation of T_iso_/2 generations. In the Isolation Migration (IM), Secondary Contact (SC) and Periodic Secondary Contact (PSC) scenarios, gene exchange was currently ongoing between the two populations. In the SC and PSC scenarios, the two populations first evolved in strict isolation and then experienced a period of gene exchange (T_sc_). In the SC scenario, there was a single period of recent migration whereas in the PSC scenario a period of ancient migration also occurred after 1-T_sc_/2 generations of strict isolation. The last scenario is the standard isolation with migration (IM) scenario in which migration occurred continuously over time since the two species started to diverge. For models including migration (IM, AM, PAM, SC and PSC), we compared two alternative models in which the effective migration rate was either homogeneous (“homo”) or heterogeneous (“hetero”) among loci (Roux *et al*. 2013, 2014, 2016). These models aim to account for the effects of a semi-permeable barrier to gene flow: a reduced effective migration rate is predicted in chromosomal regions linked to incompatible genes; while free introgression is predicted in loosely linked regions.

#### Coalescent simulations

For each of the five sampling strategies, we performed one million multilocus simulations under the eleven scenarios of speciation using the coalescent simulator Msnsam (Hudson 2002; Ross-Ibarra *et al*. 2008). Simulations fitted to the characteristics of each data set (“exome capture” data with *n* = 2, 4, 8 individuals and “rna-seq” data with *n* = 2, 4 individuals). We assumed free recombination between contigs, and we fixed the intra-contig population recombination rate to be equal to the population mutation rate. Previous studies have shown that methods which take intra-locus recombination into account remain valid when rates of recombination are low (Becquet & Przeworski 2007). Moreover, our method does not rely on haplotypic data, and so not estimating exact rates of recombination should not affect our results. To account for errors in identifying the ancestral allele in the unfolded jSFS, we explicitly modeled a misorientation rate in our coalescent simulations. We assumed that a proportion *e* of SNPs, which was a parameter to be inferred, were misoriented and changed *e*_i_, their frequency in population i, into 1-*e_i_*.

Prior distributions for *θ_A_*/*θ_ref_*, *θ_1_*/*θ_ref_* and *θ_2_*/*θ_ref_* were uniform on the interval 0–20 with 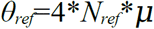. The effective size of the reference population, *N_ref_*, used in coalescent simulations was arbitrarily fixed to 100,000. The mutation rate, *μ*, was set to 2.763*10^−8^ per bp per generation (Roux *et al*. 2014). The *T_split_*/4*N_ref_* ratio was sampled from the interval 0-25 generations, conditioning the parameters *T_iso_* and *T_sc_* to be uniformly chosen within the 0-*T_split_* interval. For the scenarios including migration, we used the scaled effective migration rates *M*=4*Nm*, where *m* is the fraction of the population made up of effective migrants from the other population at each generation. In the homogeneous model, a single effective migration rate, shared by all loci but differing in each direction of introgression, was sampled from an uniform distribution in the interval 0-40. For the heterogeneous model, we assumed two categories of loci occurring in proportion *p* and (1-*p*). The parameter *p* was sampled from an uniform distribution in the interval 0-1. The first category of loci are neutral genes introgressing at a migration rate sampled from an uniform distribution in the interval 0-40. The second category comprises loci affected by the barrier to gene flow, so that their effective migration rate was reduced by a gene flow factor, *gff*, compared to neutral loci (Barton & Bengtsson 1986). The *gff* was sampled from a Beta distribution with two shape parameters (*alpha* chosen in the interval 0.001-10 and *beta* chosen in the interval 0.001-5). Prior distributions were computed using a modified version of the program Priorgen (Ross-Ibarra *et al*. 2008) as described in Roux *et al*. (2013).

#### Model choice

To choose the best supported model, we followed the methods previously described in Roux *et al*. (2013, 2014). Briefly, posterior probabilities for each of the eleven speciation scenario were estimated with a neural network using the R package abc (Csilléry *et al*. 2012). It implements a nonlinear multivariate regression by considering the model itself as an additional parameter to be inferred. The 0.01% replicate simulations nearest to the observed values of the summary statistics were selected. Moreover, to evaluate the relative probability of the heterogeneous model of migration rates across loci, we compared the alternative models (“homo” vs. “hetero”) within each scenario including gene exchange. We approximated Bayes Factors, *BF*, as the posterior probability of the best supported model divided by that of the model with the highest posterior probability from the remaining candidates. The posterior probability of each model calculated among the eleven models of speciation are detailed in Table S1; and those of the best model between heterogeneous and homogeneous migration rates are given in Table S2.

#### Model checking

We checked our ability to correctly recover the true model under our ABC framework by a “leave-one-out cross validation” from our simulations. We randomly extracted 100 simulated datasets from the million simulations performed for each model. For each of the 100 of datasets simulated under a given model, we applied the model choice procedure described above to compute the posterior probability of all competitive models. The accuracy rate for model *M* was calculated as the proportion, among simulated data inferred to correspond to model *M*, of those actually generated under model *M*. The ambiguity rate was computed as the proportion of simulated data generated under model *M* whose best model was not strongly supported, i.e. its posterior probability was below an arbitrary threshold (P_min_) set to be one third above the expected value given the total number of models compared (1/11 for eleven models, 1/6 for six models or 1/2 for two models). The accuracy and ambiguity rates for the “exome capture” data with *n*=2 individuals are provided in Table S3.

## Results

### Patterns of polymorphism

We obtained polymorphism data for two mussels species, *M. edulis* and *M. galloprovincialis*, and one outgroup (*M. trossulus*) in samples of increasing size (*n*=2, *n*=4 and *n*=8) and for the two datasets (“exome capture” vs. “rna-seq”). The jSFS of each data set is shown in Figure S1, while Table 2 gives summary statistics of genetic polymorphism.

The data produced by the two sequencing techniques differed in the total number of SNPs (*n*_snp_=3,993 (*n*=2); n_snp_=5092 (*n*=4); *n*_snp_=5000 (*n*=8) in the “exome capture” data; *n*_snp_=17,275 (*n*=2); *n*_snp_=17,902 (*n*=4) in the “rna-seq” data). The substantially lower number of SNPs in the “exome capture” data reflects the lower number of loci retained for the analysis due to a reduced and more heterogeneous sequencing depth compared to the “rna-seq” data (while applying identical coverage filters to call SNPs). However, the jSFS calculated from the two techniques had a similar proportion of sites in the different classes (Pearson’s correlation between jSFS: R^2^=0.97, p<0.0001 for *n*=2 individuals; and R^2^=0.99, p<0.0001 for *n*=4 individuals). Globally, the mean level of genetic differentiation was quite low (FST~10% in the “exome capture” data; FST~18% in the “rna-seq” data, see Table 2), but highly variable across loci (standard deviation was ~16 % and ~24 %, respectively). A low proportion of sites (<5% in the “exome capture” data and <10% in the “rna-seq” data) were fixed differences, while two to three times more SNPs were polymorphic and shared between the two species (see Figure S1). The level of intraspecific nucleotide diversity was elevated (π_edu_=π_gal_=0.016 in the “exome capture” data; π_edu_=0.038 and π_gal_=0.036 in the “rna-seq” data for *n*=2, Table 2) and not significantly different between the two species (non-significant Wilcoxon signed-rank test). Polymorphic sites were mainly private to each species (~80% of the sites), and mainly corresponded to low frequency classes. Moreover, the jSFS was remarkably symmetric suggesting limited differences in population size and/or migration rates between the two species. These patterns were consistent across sample sizes, but there were some differences comparing the two techniques. Specifically, the “exome capture” data set showed significantly lower level of divergence (Wilcoxon signed-rank test, p<0.0001 between netdiv_capture_=0.004 and netdiv_rna-seq_=0.02 for *n*=2 individuals, Table 2) and average number of fixed differences between species per locus (Wilcoxon signed-rank test, p<0.0001 between Sf_capture_=0.322 and Sf_rna-seq_=0.810 for *n*=2 individuals, Table 2). These discrepancies were most likely due to the use of a single reference in the “exome capture” data resulting in the problematic mapping of highly divergent alleles from the two species.

### Effects of the number of individuals and SNPs on model selection

We carried out model selection between the various scenarios of speciation shown in Figure 1, and asked whether the number of individuals and number of SNPs had an effect. Results are shown in Table 3a, which reports the posterior probability of the best supported scenario for each sampling strategy; and in Table 3b which compares the posterior probability of homogeneous *vs*. heterogeneous migration for the best supported scenario (see also Table S1 and S2 for full details).

**Table 3.**
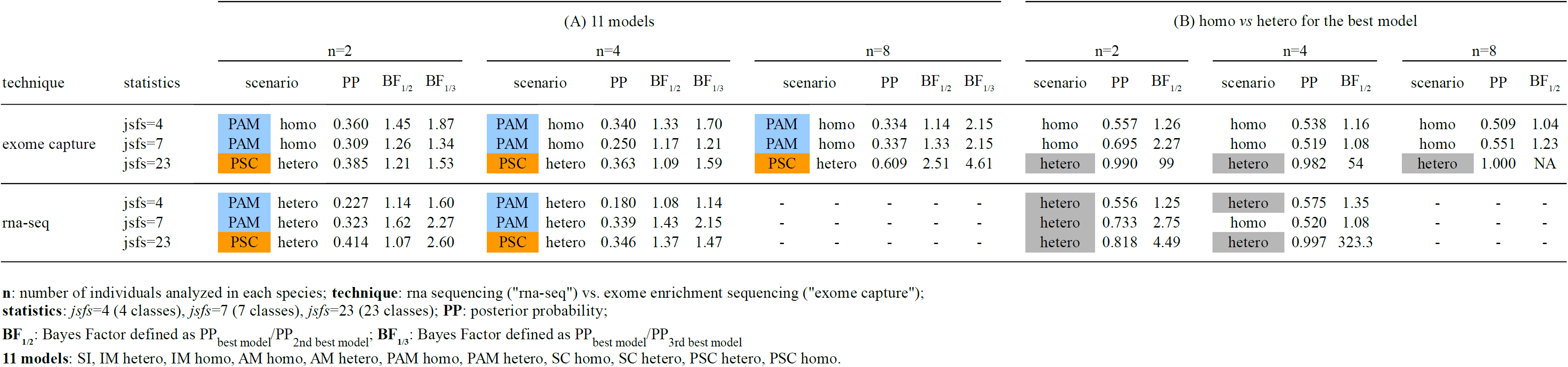
Posterior probabilities of the speciation models

Firstly, we compared the “exome capture” and the “rna-seq” data which differ in the number of SNPs sampled (3,993 SNPs and 17,275 SNPs, respectively); and we found that the best supported scenario was the same for both data sets (Table 3a). For example, when considering twenty-three classes in the jSFS (*jsfs*=23), the best supported scenario always involved recent and genome-wide heterogeneous migration between the two species. The heterogeneous periodic secondary contact (PSC.hetero) was the most supported scenario; and the next best model described a very similar history of secondary contact with a single period of gene exchange: *BF*=P_PSC.hetero_/P_SC.hetero_=1.21 with the “exome capture” data, *BF*=P_PSC.hetero_/P_SC.hetero_=1.07 with the “rna-seq” data, in the case of *n*=2 individuals. With *n*=4 individuals, for which the next best model also included gene flow, those numbers were P_PSC.hetero_/P_IM.hetero_=1.09 and P_PSC.hetero_/P_IM.hetero_=1.37. The same patterns were found when using different subsets of the jSFS (*jsfs*=4 and *jsfs*=8, Table 3a), except that the best supported scenario was then the periodic ancient migration (PAM). Regarding genome-wide heterogeneity of migration rates (Table 3b), the “rna-seq” data gave more support to the heterogeneous model (e.g., *BF*=P_AM.hetero_/P_AM.homo_=1.25 with *n*=2 and *jsfs*=4) compared to the “exome capture” data (*BF*=P_AM.homo_/P_AM.hetero_=1.26). This is consistent with the higher heterogeneity of the jSFS in the “rna-seq” data, involving a higher proportion of fixed differences and shared polymorphic sites (Table 2 and Figure S1).

Secondly, we evaluated the effect of the number of individuals sampled. As with the number of SNPs, it is clear that sample size had little effect. The best supported scenario remained consistent across the different sampling size (*n*=2, 4 or 8 individuals; Table 3a). For example, when considering twenty-three classes in the jSFS (*jsfs*=23), the heterogeneous periodic secondary contact scenario was the best supported model whatever the sampling size in both datasets.

### Effects on model selection of the number of classes in the jSFS

We next investigated the effects of binning the jSFS: (i) on model choice in the mussel datasets (Figure 2 and Table 3), and (ii) on our ability to discriminate between different speciation scenarios based on simulated datasets (Table S3). Given the limited effects of the sampling strategy, the ABC performance results are presented for a simulated dataset resembling the “exome capture” data with *n*=2 individuals only. This allowed us consider the full spectrum when using twenty-three classes of polymorphism.

#### Heterogeneity of migration rates

The scenarios with recent migration (SC, PSC and IM) all strongly supported heterogeneity of migration rates; and this support tended to increase with the number of polymorphic classes considered (Table S2). For example in the “exome capture” data with *n*=2 individuals, the relative probability of the heterogeneous model in the periodic secondary scenario was P_PSC.hetero_=0.53 with *jsfs*=4, P_PSC.hetero_=0.77 with *jsfs*=7 and P_PSC.hetero_=0.99 with *jsfs*=23. In contrast, the model of homogeneous migration rates was the most supported, though marginally, in the scenarios of ancient migration (PAM and AM): e.g., P_PAM.homo_=0.56 with *jsfs*=4, P_PAM.homo_=0.69 with *jsfs*=7 and P_PAM.homo_=0.63 with *jsfs*=23. Concordant patterns were obtained using the other spectra, which differed in the number of individuals and SNPs sampled (Table S2).

Using simulated datasets, we then assessed the performance of the method in identifying the correct model when homogeneous and heterogeneous models were compared (Table S3a). The correct model was always recovered (i.e., an accuracy rate of 1) for the different binnings in all speciation scenarios including gene flow; however, the ambiguity rate did strongly decrease when more information from the jSFS was included. With only four classes (*jsfs*=4), none of the replicates showed a posterior probability higher than 0.83 (the threshold set for the 2-model comparison), which corresponds to an ambiguity rate of 1. Similarly, all ambiguity rates were above 0.97 with seven classes (*jsfs*=7). In contrast, when considering the full jSFS (*jsfs*=23), we could correctly recover with a strong support the simulated speciation models (e.g., 63% of the PSC.hetero replicates and 43% of the PAM.hetero replicates were above the threshold, Table S3a). These results suggest that the additional classes of the 23-binned jSFS are necessary to detect heterogeneity of migration rates across the genome.

#### Scenarios of speciation

It is clear from Table 3a (see details in Table S1) that binning the jSFS to four or seven classes leads to a loss of information. Specifically, when considering the full jSFS (*jsfs*=23), only the scenarios involving recent migration were supported (P_PSC_+P_SC_+P_IM_=0.96, Table S1); while the contrary was true when fewer classes of polymorphism were used (P_PAM_+P_AM_=0.96 with *jsfs*=4 and 0.93 with *jsfs*=7, Table S1). Remarkably the strict isolation scenario was never supported (P_SI_<0.06, Table S1) suggesting that gene flow must have occurred between the two mussels species during divergence. Moreover, the fact that the most supported scenario, using full information (*jsfs*=23), was the heterogeneous periodic secondary contact (P_PSC.hetero_=0.39) suggests a complex history of speciation, including periods of isolation alternating with ancient and recent migrations. The discrepancies that appear when not distinguishing low and high frequency shared variants (in the case of *jsfs*=4 and *jsfs*=7), confirm their importance for the identification of recent migration events (Alcala *et al*. 2016). In general, the best supported model fits well the data for each binning strategy (Figure S2). All observed statistics were in the 95% simulated posterior distribution, except for the class “ssfB_2” (derived polymorphism fixed in species B, but polymorphic (doubletons) in species A) which was slightly overestimated by the model PSC.hetero (*jsfs*=23, Figure S2a); and the class “sfB” (derived polymorphism fixed in species B, and absent in species A) which was slightly underestimated by the model PAM.hetero (*jsfs*=7, Figure S2b). No statistics were found significantly out of the simulated distribution under model PAM.hetero with *jsfs*=4 (Figure S2c). On the contrary, the alternative models failed to fit more parts of the spectra. For example with *jsfs*=23 (Figure S2a), in which PSC.hetero is the best model (P_PSC_=0.385), the alternative model PAM.hetero (P_PAM_=0.007) overestimates the number of derived sites fixed within each species in all ‘sf’ classes and model SI (P_SI_=0.009) underestimates the number of shared polymorphisms in most ‘ss’ classes.

We further evaluate, by simulation, the effect of binning the jSFS on our capacity to infer the correct speciation model (Table S3b). As the heterogeneous models consistently outperformed the homogeneous models in scenarios with ongoing migration, and they were not significantly less likely in the models of ancient migration (Table S2), the ABC performance was evaluated among the five models of migration including heterogeneous migration only (IM.hetero, SC.hetero, PSC.hetero, AM.hetero and PAM.hetero), plus the strict isolation model (SI). Globally, the probability of rejecting the correct model decreased when increasing the number of polymorphic classes. By using four or seven classes *vs*. twenty-three classes, the correct model was recovered at an estimated rate equal to or lower than 0.70 *vs*. 1 for IM.hetero, 0.58 *vs*. 0.50 for SC.hetero, 0.89 *vs*. 1 for PSC.hetero, 0.34 *vs*. 0.42 for AM.hetero, 0.70 *vs*. 0.79 for PAM.hetero and 0.73 *vs*. 0.84 for SI (Table S3b). Moreover, we could not discriminate between the different scenarios of recent migration (PSC.hetero, SC.hetero and IM.hetero) when using only four or seven classes (Figure S3a); and the same was true with the two scenarios of ancient migration (PAM.hetero and AM.hetero, Figure S3b). In contrast, the full jSFS (*jsfs*=23) contains enough information to more accurately identify the periodic secondary contact as the true model among the other scenarios of recent migration (Figure S3a). Nevertheless, distinguishing between the two ancient migration scenarios remained difficult even with *jsfs*=23 (Figure S3b). Finally, the ambiguity rates also strongly decreased when binning the jSFS in more classes: all ambiguity rates were equal to or greater than 0.86 in the recent migration scenarios, 0.41 in the ancient migration scenarios and 0.21 in the strict isolation scenarios with *jsfs*=4 or *jsfs*=7; whereas these numbers were 0.25, 0.20 and 0.15 with *jsfs*=23 (Table S3b).

## Discussion

NGS data give us the opportunity to capture the diversity of coalescent histories across loci, and so to reveal the complexity of the speciation process (Sousa & Hey 2013). Recently, important efforts have been made to develop statistical methods of inference making use of population genomics data. Computing the joint site frequency spectrum (jSFS) is an efficient way of summarizing the demographic information contained in NGS data because anonymous SNPs can be used (e.g. produced by RAD-sequencing) and it does not rely on phased data. However, the jSFS obtained from low-coverage sequencing data (typically <10x per position and per individual) can be biased toward a deficit of rare variants; and this is of particular concern when investigating the demographic history of populations (e.g., Nielsen *et al*. 2012; Han *et al*. 2013). A first category of maximum-likelihood methods uses forward diffusion theory to compute numerical solutions to the jSFS under complex models (see Gutenkunst *et al*. 2009; Lukic & Hey 2012). A second category of methods estimates the expected jSFS under any demographic model, as simulations are used to approximate the composite-likelihood of the data (see Naduvilezhath *et al*. 2011; Excoffier *et al*. 2013). Here, we used a likelihood-free method (Approximate Bayesian Computation) on the joint site frequency spectrum of coding sequences, and we evaluated the influence of different sampling strategies on the inference of speciation models in mussels.

Our ABC-based model comparison shows little qualitative effect of individual and SNP sampling on the outcomes. First, inferences based on *n*=2, *n*=4 or *n*=8 individuals supported the same model of periodic secondary contact (Table 3), confirming previous results that relatively few individuals are sufficient to make robust inferences on the way divergence occurs between lineages (e.g., Robinson *et al*. 2014). This is because most coalescence events occur recently in the population history; so increasing the number of individuals is only helpful to characterize recent demographic events rather than the past divergence history. Second, we showed that inferences from the two sequencing techniques (“exome capture” vs. “rna-seq”) were qualitatively the same, despite the very different number of SNPs that were sampled, again supporting the periodic secondary contact scenario (Table 3). This implies that neither the sequencing technique, nor the number of informative sites have a substantial effect on the inference. In fact, the jSFS calculated from the different data sets were very similar (Figure S1); we only found a deficit of divergent SNPs in the “exome capture” data that may be due to the difficulty of mapping highly divergent alleles onto a single reference (on the contrary, the “rna-seq” reads from the different species were independently assembled and mapped). From a theoretical perspective, adding more loci (or longer loci) provide information about deep coalescent events which are important for shedding light on the divergence history of closely-related species (e.g., Wang & Hey 2010). In fact, previous simulation studies showed an influence of locus length and locus number on the ABC performance in model choice, and highlighted a threshold effect above which adding more loci did not significantly improve inferences (e.g., Li & Jakobsson 2012; Robinson *et al*. 2014; Shafer *et al*. 2015). Thereby, we argue that the number of SNPs sampled in our study was sufficient to accurately represent the diversity of coalescent histories in the *Mytilus* genome, and consistently support the same speciation model.

In most studies, functions of the jSFS are used as summaries of the data. For example, the likelihood method of Nielsen & Wakeley (2001) uses four classes of polymorphisms to estimate migration rates and divergence times in the isolation with migration scenario. In the ABC approach, choosing a suitable set of summary statistics is difficult because it implies a trade-off between loss of information and reduction of dimensionality. Accordingly, practical methods to identify approximately sufficient statistics have been developed (e.g., Wegmann *et al*. 2009; Nunes & Balding 2010; Aeschbacher *et al*. 2012). Here, we compared ABC inferences based on 23-binned jSFS (*jsfs*=23) with inferences using a subset of polymorphic classes (*jsfs*=4 and *jsfs*=7). Our simulation results pointed out the loss of information when only four or seven classes in the spectrum were considered; particularly for the inference of recent migration events. By decomposing the jSFS into twenty-three classes, we could reveal the excess of shared polymorphisms that are at high frequency in one species and low frequency in the other, a pattern produced by recent migrants (Table 3). This is in agreement with the simulation study of Tellier *et al*. (2011) that showed a significant improvement in the estimation of the timing of gene flow when these additional polymorphic classes were considered; and the study of Alcala *et al*. (2016) that showed an excess of high frequency derived alleles is the characteristic footprint of secondary contacts. Across all models, we showed that the probability to correctly infer the true model (accuracy rate) increases with the number of classes considered, and the ambiguity rate correspondingly decreases (Table S3). Smith *et al*. (2017) similarly found that the statistical power of their ABC model selection increases with the number of classes until reaching a plateau of error rates (but above which computation efforts continues to increase).

Inferences based on the 23-bin jSFS consistently support a model of periodic secondary contact with genomic heterogeneity in gene flow (PSC.hetero) in both the “exome capture” and “rna-seq” mussel datasets, and simulations showed that this model has a high accuracy rate, and one of the lowest ambiguity rates (Table S3). Although its relative posterior probability was moderately higher than that of the other models with migration (SC and IM, Table S1), we showed that the method has some power to distinguish PSC from these models (Figure S3a). These results are in agreement with previous ABC-based studies revealing clearly a secondary contact history between the two mussel species (Fraïsse *et al*. 2014; Roux *et al*. 2014), although they relied on eight nuclear loci and on a different set of summary statistics (those that we called “mscalc” here). The periodic connectivity models (PSC and PAM) were not included in these previous studies because of the lack of power to test for intermittent gene exchange since secondary contact. In the present study, we directly compared the use of the jSFS *vs*. “mscalc” statistics by performing additional ABC inferences on the “mscalc” statistics presented in Table 2. Results were similar to those obtained with *jsfs*=23 (Table S1), i.e. the best supported models included one or two secondary contacts, except for two datasets (“exome capture” data with *n*=2 and *n*=8 individuals) for which strict isolation was chosen with a posterior probability of 37% and 34%, respectively. However, the goodness-of-fit of the strict isolation model was quite poor for the standard deviation of the number of fixed sites, “sf_std”, and very poor for the standard deviation of the FST, ‘fst_std’ (“exome capture” data with *n*=2 individuals, Figure S2d). Both statistics were underestimated by the model suggesting that a history without gene flow cannot produce the observed variation of genetic divergence along the genome. Across models, inferences based on the “mscalc” statistics showed similar accuracy rate to those based on *jsfs*=23; however, the ambiguity rates for models with current migration were somewhat higher (PSC = 0.52 *vs*. 0.30, SC = 0.61 *vs*. 0.31, IM = 0.40 *vs*. 0.25, respectively). These supplementary analyses suggest that extracting summary statistics from the jSFS can lead to a substantial loss of information.

## Conclusion

In this work, we have shown that two high-throughput sequencing datasets (“exome capture” and “rna-seq”), imply the same history of divergence in mussels, regardless of the number of individuals or SNPs sampled, but conditional on the inclusion of informative classes in the joint site frequency spectrum. Thus, genome-wide data coupled with flexible inference methods allow us to test for more complex scenarios of divergence, by providing a comprehensive picture of the gene histories across the genome. Here, we incorporate in our ABC framework heterogeneity in migration rates among loci, to account for the semi-permeability of the barrier to gene flow between recently diverged species (Barton & Bengtsson 1986). This variation in the rate of effective migration results in variation of genetic divergence along the genome. As shown elsewhere (Roux *et al*. 2013, 2014, 2016), failing to account for this heterogeneity can mislead inferences. In a similar way, the effect of background selection (i.e. purifying selection at linked loci) and genetic hitchhiking (i.e. positive selection at linked loci) in regions of low recombination can now be incorporated in demographic inferences by including heterogeneity in effective sizes among loci (Sousa & Hey 2013, Roux *et al*. 2016, Aeschbacher *et al*. 2017). Combining modelling approaches that account for both sources of genomic heterogeneity (e.g., Roux *et al*. 2016) may provide further insight into the complex interplay between linked selection and resistance to introgression during speciation with gene flow (e.g., Duranton *et al*. 2018).

## Acknowledgements

The computations were performed at the “Montpellier Bioinformatics Biodiversity” computing cluster platform (CeMEB LabEx “Mediterranean Center for Environment and Biodiversity”). This work was funded by the Agence Nationale de la Recherche (HYSEA project, ANR-12-BSV7-0011) and by a Languedoc-Roussillon Region “Chercheur(se)s d’avenir” grant (Connect7 project). This is article XXX of “Institut des Sciences de l’Evolution de Montpellier”.

## Data Accessibility

**Data set 1: “exome capture”:** http://www.scbi.uma.es/mytilus/index.php (Fraïsse *et al*. 2016).

**Data set 2: “rna-seq”:** http://kimura.univ-montp2.fr/PopPhyl (Romiguier *et al*. 2014).

## Supplementary Figures

**Figure S1.** Unfolded joint site frequency spectrum for the two different sequencing techniques.

**Figure S2.** Goodness-of-fit test for three models (PSC, PAM and SI) with the “exome capture” data, *n*=2 individuals and the different summary statistics.

**Figure S3.** Evaluation of the ABC performance on model choice.

## Supplementary Tables

**Table S1.** Posterior probabilities - 11 models.

**Table S2.** Posterior probabilities - homo vs. hetero.

**Table S3.** Accuracy and ambiguity rate (*n*=2, “exome capture” data).

## Supplementary Methods

**Text S1.** Scripts to compute summary statistics from the data. **a**. Extracts polymorphic synonymous positions from a fasta alignment. **b**. Produces the joint site frequency spectrum for each contig, and mscalc statistics across contigs. **c**. Produces a summed jSFS across contigs.

**Text S2.** Scripts to compute summary statistics from simulations. **a**. Produces coalescent simulations under a set of demographic models, and compute joint site frequency spectrum for each contig, and mscalc statistics across contigs. b. Produces a summed jSFS across contigs.

**Text S3.** Scripts to estimate the posterior probability of each demographic model using neural networks based on jSFS (**a**) and mscalc statistics (**b**). Scripts to evaluate the ability of the method to correctly recover the true simulated model based on jSFS (**c**) and mscalc statistics (**d**).

